# Autotaxin Inhibition Ameliorates HFpEF Phenotype By Reducing LPA-Mediated Systemic Inflammation and Cardiac Remodeling

**DOI:** 10.64898/2026.08.26.747366

**Authors:** Rajesh Chaudhary, Andrew Robbins, Anand P. Singh, Parisa Sabahni, Tahra K Luther, Afnan Alzamrooni, Rachel Lopez, Tania Maheshwari, Nicolas Collins, Scott Hummel, Ahmed Abdel-latif

## Abstract

**Background:** HFpEF accounts for roughly half of heart failure admissions and lacks disease-modifying therapy. Autotaxin (ENPP2) generates lysophosphatidic acid (LPA), a profibrotic and pro-inflammatory bioactive lipid. Whether circulating lysophospholipid metabolism is altered in HFpEF, and whether autotaxin inhibition modifies an established experimental HFpEF phenotype, is untested.

**Methods:** Plasma from patients with HFpEF (n=210) and non-heart-failure comparators (n=27) underwent untargeted and LPA-targeted mass spectrometry and a nine-analyte multiplex immunoassay. Male C57BL/6J mice received a high-fat diet plus L-NAME (0.85 g/L) or chow for 5 weeks; after phenotype confirmation, they received oral PF-8380 (30 mg/kg/day) or vehicle for 10 weeks. Endpoints were echocardiography, functional assessment, gravimetric studies, tail-cuff pressure, trichrome fibrosis, and flow cytometry of heart and spleen.

**Results:** All nine analytes, including the autotaxin protein ENPP2, were higher in HFpEF than comparators. HFpEF plasma showed higher LPE O-16:1, LPE O-18:2, PS 38:4 and PC 36:4;O, and lower SM 39:2;O3 and PS 36:0. LPA 20:0 was 3.5-fold higher in both sexes, whereas LPA 18:2 was lower in women. Diet plus L-NAME raised blood pressure, LV mass, and isovolumic relaxation time with preserved ejection fraction. PF-8380 reduced echocardiographic indices of diastolic dysfunction, fibrosis area, cardiomyocyte area, and cardiac CD11b+, CD64+, CD86+, and Ly6G+ frequencies, without a change in fat or lean mass.

**Conclusion:** In male mice with established two-hit HFpEF, autotaxin inhibition improved diastolic indices and reduced fibrosis, hypertrophy and cardiac myeloid accumulation. Human data show altered lysophospholipid composition. Collectively, these findings nominate the autotaxin–LPA axis as a tractable therapeutic target and support further evaluation of autotaxin inhibition as a candidate disease-modifying strategy for HFpEF management.

**Graphical Abstract:** 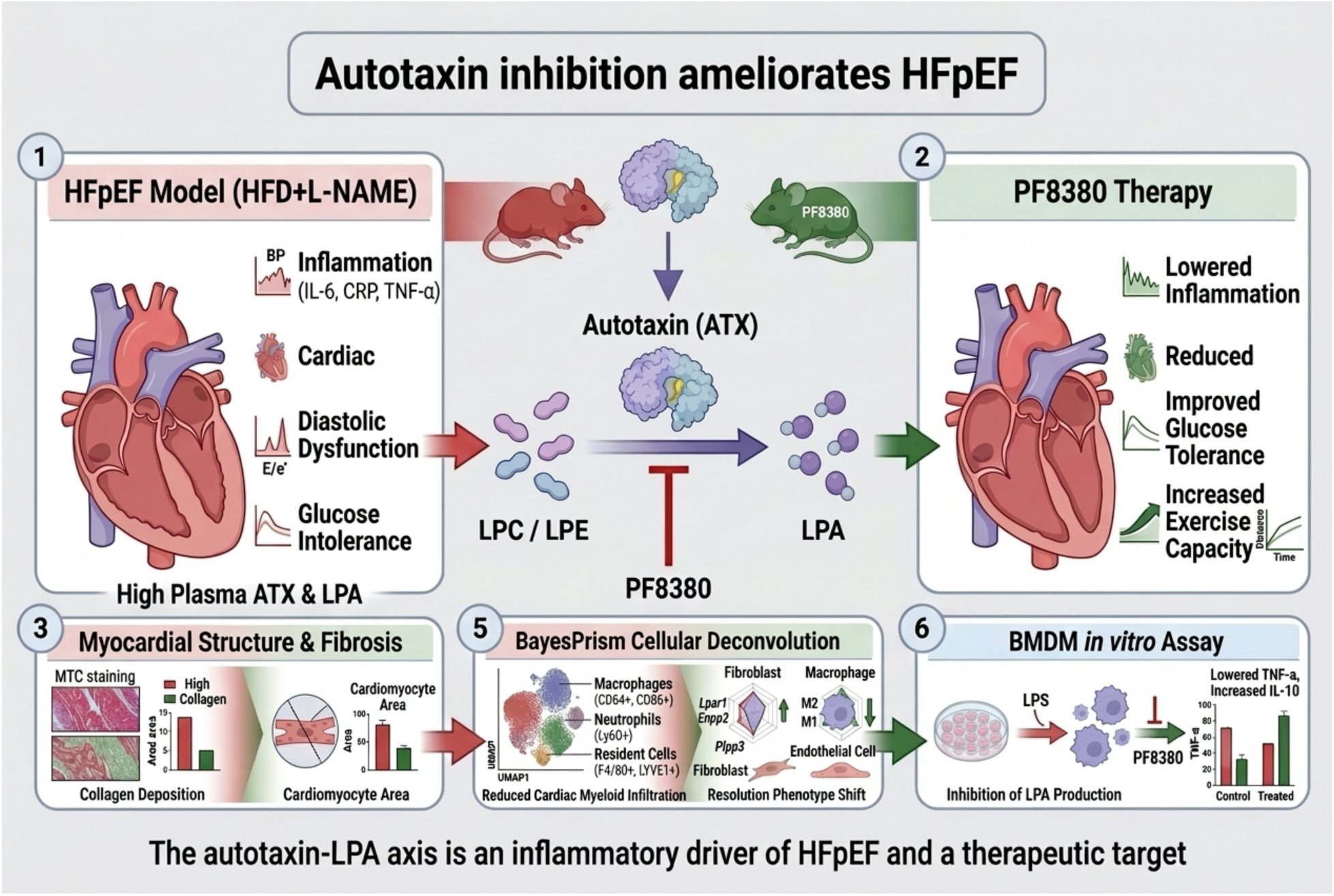

## Introduction

Heart failure with preserved ejection fraction (HFpEF) is now the most common form of heart failure, yet it remains a condition with markedly limited treatment options. HFpEF accounts for approximately half of all heart failure and is its fastest-growing phenotype, driven by the rising prevalence of obesity, hypertension, and type 2 diabetes in an aging population (1). Five-year mortality exceeds 50% and approaches that of heart failure with reduced ejection fraction, yet therapeutic options remain sparse. Sodium-glucose cotransporter 2 inhibitors (SGLT2i) and glucagon-like peptide-1 (GLP1) receptor agonists reduce heart failure hospitalization but have not established a clear survival benefit, and no available therapy reverses the myocardial remodeling that underlies the syndrome (2, 3). This gap reflects an incomplete understanding of the mechanisms that link systemic metabolic stress to diastolic dysfunction, cardiac fibrosis, and exercise intolerance frequently observed in HFpEF.

A common pathology shared across the comorbidities that precipitate HFpEF is chronic, low-grade systemic inflammation (4). Obesity, hypertension, and diabetes generate a sterile inflammatory state that promotes coronary microvascular endothelial dysfunction, myocardial stiffening, and interstitial fibrosis. Circulating inflammatory markers, including interleukin-6 (IL-6), growth differentiation factor-15 (GDF-15), C-reactive protein (CRP), and tumor necrosis factor-alpha (TNF-α), are elevated in patients with HFpEF and track with the severity of diastolic dysfunction (5–7). The upstream mediators that initiate and sustain this response, and whether they can be targeted pharmacologically, remain poorly defined. Bioactive lipids are compelling candidates because they couple the metabolic derangements of HFpEF to inflammatory and fibrotic signaling, and their accumulation in the failing myocardium is associated with adverse remodeling(8).

Among bioactive lipids, lysophosphatidic acid (LPA) is a potent regulator of inflammation and tissue fibrosis. LPA is generated extracellularly by autotaxin (ENPP2, ectonucleotide pyrophosphatase/phosphodiesterase 2), a secreted lysophospholipase D that hydrolyzes lysophosphatidylcholine and lysophosphatidylethanolamine into LPA, which then signals through six G-protein-coupled receptors (LPA1 to LPA6) (9). The autotaxin-LPA axis recruits and activates myeloid cells (Zhao et al., 2015) and drives the fibroblast-to-myofibroblast transition that produces organ fibrosis. Genetic or pharmacologic disruption of this axis attenuates fibrosis in the lung (10, 11) kidney (12), and heart (13, 14), and we and others have shown that autotaxin inhibition reduces myocardial inflammation and adverse remodeling after ischemic injury (15–17). LPA receptor 1 antagonism has advanced to clinical testing in idiopathic pulmonary fibrosis (18). Despite this rationale, whether the autotaxin-LPA axis is engaged in HFpEF and whether its inhibition is therapeutic have not been tested.

We hypothesized that the autotaxin-LPA axis is activated in HFpEF and drives the systemic inflammation and cardiac remodeling that define the syndrome. To test this, we paired targeted and untargeted plasma lipidomics with multiplex inflammatory profiling in patients with HFpEF and controls, and we inhibited autotaxin with the small-molecule inhibitor PF8380 in a two-hit (high-fat diet plus L-NAME) mouse model of HFpEF (19). To resolve the cellular basis of autotaxin inhibition, we integrated high-dimensional spectral flow cytometry and functional assays in bone marrow-derived macrophages. Our results identify the autotaxin-LPA axis as an inflammatory driver of HFpEF and establish that its inhibition is a therapeutic strategy that reduces systemic inflammation, cardiac myeloid cell accumulation, and adverse cardiac remodeling.

## Study design and methods

### Human samples

The University of Michigan Medicine School Central Biorepository (UM CRB) obtained plasma and serum samples under the oversight of the University of Michigan School Institutional Review Board (IRB). Participants were enrolled in the UM CBR during clinical encounters at Michigan Medicine after they were informed about the study’s purpose and design, and they signed informed consent for the collection, storage, and future research use of their biological samples. Participants could withdraw from the study at any time. HFpEF was defined as signs and symptoms of heart failure (HF) with left ventricular ejection fraction (EF) (LVEF) ≥50%, as previously described (20). We followed the inclusion and exclusion criteria for the study group as previously defined (20). We collected and processed total blood to separate plasma and serum, then stored it at-80 °C until further processing, following the standardized institutional procedure. Trained biorepository personnel processed samples using a harmonized operating procedure to ensure consistency across participants. LPA can be generated or degraded after blood collection, and platelet activation can substantially influence individual species. Therefore, trained biorepository personnel carefully handled samples, and the downstream process was managed to minimize processing time and temperature, including freeze–thaw cycles. Clinical metadata – including demographics, diagnoses, laboratory values, and relevant comorbidities- were curated and linked to the biospecimens through the biorepository’s secure data infrastructure. For this study, the UM CBR released de-identified aliquots of plasma and serum from 237 participants: 27 controls and 210 with heart failure with preserved ejection fraction (HFpEF).

## Animal study

### Research design and methods

#### 2.1 Animals

All animal experiments followed the approved University of Michigan Institutional Animal Care and Use Committee (IACUC) protocol. Eight-week-old male C57BL/6 mice of J strains (n=10-12/group, Jackson Laboratory, Bar Harbor, ME) were included in this study and were randomized to one of the four groups: 1) Chow+Vehicle, 2) Chow+PF8380, 3) HFD+L-NAME+Vehicle, 4) HFD+L-NAME+PF8380, after undergoing two-week acclimatization at an ambient temperature of 23 °C prior. We previously showed that housing temperature alters the metabolic phenotype of this HFpEF model in C57BL/6J mice (Chaudhary et al., 2025). All mice were housed in 12h of light:12h of darkness with lights off at 07:00 (Zeitgeber, ZT 0) and lights on at ZT 12 and received either Chow (18% calorie from fat) and/or high-fat diet (HFD, #D12492, Research Diet, USA, 60% of calorie from fat) in combination with nitric oxide synthase inhibitor, *N^w^*-nitro-L-arginine methyl ester (L-NAME, 0.85g/L (#32870050, Thermo Scientific, USA) in drinking water. All mouse groups received food and water ad libitum throughout the 15-week study. We changed food, cages, and water twice weekly for all cages, including the Chow group, to standardize handling, and recorded body weights weekly during cage changes at ZT14. Parts of heart, liver, spleen, whole blood, and total bone marrow were snap-frozen in liquid nitrogen and stored at -80 °C for further processing. We perfused the heart with cold phosphate-buffered saline and separated it for bulk RNA sequencing, flow cytometry, and histology. Whole blood was collected via direct cardiac puncture, placed in tubes containing CTAD at 1:10, and plasma samples were prepared after spinning at 700g for 5 min at room temperature. The supernatant was collected and centrifuged again at 10,000g for 10 min at room temperature.

#### 2.2. Intraperitoneal glucose tolerance test

Intraperitoneal glucose tolerance (IPGTT) was performed at week 5 and 15 following 5h of fasting using a bolus dose of 1g/kg body weight (n=7-10 mice/group). Blood glucose readings were measured at 0, 15, 30, 60, 90, and 120 min following the bolus dose via tail-vein prick using Accu-Chek® Performa II, Roche, USA. The area under the curve (AUC) was calculated using the trapezoid rule (21).

#### 2.3 Body weight and composition

Body weight was recorded weekly during cage, food, and water change at ZT14. Body composition was measured in a subset of mice (n=4/group) using an NMR-based analyzer, the EchoMRI^TM^-500 body composition analyzer, according to the manufacturer’s instructions.

#### 2.4. Blood pressure measurements

Blood pressure measurements were conducted via tail cuff method using a CODA Monitor (Kent Scientific), according to the manufacturer’s instructions, after the mice were acclimatized and settled in a quiet procedure room with no disturbance, as previously described (22). Briefly, an occlusion (O) trail cuff and a volume pressure recording (VPR) cuff were placed on the tail of the mouse, and the O-cuff was inflated to impede blood flow to the tail before the cuff was deflated slowly and return blood flow was measured using a VPR sensor. Measurements were recorded after ensuring the body temperature was between 32 0C and ≤37 0C. Body temperature was recorded at the base of the tail using a laser temperature gun. All measurements were recorded in non-anesthetized mice and body temperature was maintained using a thermal pad.

#### 2.5. Echocardiography measurements

All echocardiography measurements were recorded in anesthetized mice maintained with 1-2% isoflurane and 95% oxygen to maintain the heart rate at 450 +/- 50 beats± per minute, as previously described (22). Briefly, trans-thoracic echocardiography was conducted using the Vevo F2 Imaging System (#53699-20, FUJIFILM VisualSonice, Inc., USA). The anterior chest hair was removed using Nair Hair-removal cream before sedation. Body temperature-maintained pre-warmed ultrasound gel was applied to the area underlying the heart after immobilizing the mice on the stage. The parasternal short-axis view, indicated by the presence of papillary muscle was used to obtain the M mode for ejection fraction, fractional shortening, left ventricular mass, cardiac output, and other cardiac parameters. The apical four chamber view was used to obtain the tissue Doppler and mitral valve pulse-wave Doppler measurements for myocardial tissue and blood flow velocity, respectively. All parameters were measured at least 3-5 times, and the mean was recorded as the final measurement for statistical analysis.

#### 2.6. Exercise exhaustion test

An exercise exhaustion test was performed on a subset of mice from each group, as previously described (19). Briefly, mice were acclimatized to the treadmill for 3 days prior to the experiment. Mice ran at (20 °) on a treadmill starting at 5 m/min for 4 min, and the speed was increased by 2 m/min every 2 min until the mouse was exhausted. Exhaustion was defined as the mouse’s inability to resume running within 10 sec after exposure to the electric-stimulus grid or after coming into contact with the electric-stimulus grid at least 3 times within 10 sec. Running time and distance were recorded.

#### 2.7. Histopathology

Histopathology services were performed by the In Vivo Animal Core Histology Laboratory within the Unit for Laboratory Animal Medicine at the University of Michigan and as previously described (22). Briefly, tissue was fixed in 4% paraformaldehyde (PFA) for 24h before transferring it to 70% ethanol. Fixed tissues were embedded in paraffin and sectioned at 4 μm thickness on a rotary microtome (TissueTek VIP5 ®, Sakura Finetek, USA, Inc, Torrance CA).

##### 2.7.1. Masson Trichrome

Following deparaffinization and hydration with xylene and graded alcohols, formalin-fixed, paraffin-embedded (FFPE) slides were placed in 60 °C Bouin’s Fluid (Rowley Biochemical; #F-367-1), a mordant, for 1 h. Slides were cooled briefly and then rinsed well in water. Slides were stained in Biebrich Scarlet-Acid Fuchsin (Rowley Biochemical; #F-367-3) for 10 min then briefly rinsed in deionized water before being placed in Phosphomolybdic-Phosphotungstic Acid Solution (Rowley Biochemical; #F-367-4), a mordant, for 15 min. Finally, slides were transferred directly into Aniline Blue Solution (Rowley Biochemical; #F-367-5) for 8 min, followed by a brief rinse in deionized water. Slides were dehydrated and cleared through graded alcohols and xylene and were cover-slipped with Micromount (Leica Biosystems; #3801731) using a Leica CV5030 automatic cover slipper.

#### 2.8. RNA extraction and RT-qPCR

Frozen heart tissue (20-30 mg) was pulverized at 50 Hz with TissueLyser LT (Qiagen, Hilden, Germany) and total RNA was extracted using RNeasy Mini Kit (#74106, Qiagen, Germany) as per manufacturer’s protocol. The concentration and purity of total RNA was confirmed using NanoDrop 2000 Spectrophotometer (Thermo Scientific, USA).

#### 2.9. Flow Cytometry Analysis

Flow cytometry was performed at week 15, at the end of the study, following metabolic phenotyping. Heart samples were prepared for flow cytometry as previously described (23). Briefly, mice were sedated with isoflurane before sacrificing via cervical dislocation, as per Institutional Animal Care and Use Committee (IACUC) guidelines. Hearts were perfused with cold PBS and samples were collected on cold ice before gently mincing with razor blades. Enzymatic digestion was performed by incubating the tissue samples at 37 ^0^C for 30 min in a cocktail of Collagenase type I and type XI (Sigma, #C0130 and C7657) and DNase I (Sigma, #D5307) on a rocker with gentle agitation. Cold staining buffer was used to stop the enzymatic digestion reaction, and the suspension was placed on ice. Samples were filter using 70 µm cell strainers before centrifuging at 400g for 5 min at 4 ^0^C. Samples were processed for debris and RBC removal, and cell pellets were resuspended in 1 ml of staining buffer before cells were counted using automated cell counter (Luna II, Logos Biosystems Inc., South Korea). Approximately 1 million cells were stained with fluorescently conjugated antibodies targeting macrophage-associated markers Iba1, PE-Cy5-conjugated CD64 (BioLegend, #139332), BUV615-conjugated CD11b (BD Bioscience, #751140, RRID:AB_2875166)) and validated markers distinguishing resident macrophages (CD68+ (BV711-conjugated CD68 (BioLegend, #137029, RRID:AB_2783098/CSF1R+ (BV421-conjugated CD115 (BioLegend, #135513, RRID:AB_2562667)/CD206+ (PE-eF610-conjugated CD206 (eBioscience, Thermo Fisher Scientific, #61-2061-82, RRID:AB_2802389)CD14-/IL-1β - (eF450-conjugated IL-1β, eBioscience, ThermoFisher Scientific, #48-7114-80, RRID:AB_2574108)/S100A8-) from circulating macrophages (CD14+/IL-1β+/S100A8+/CD68-/CSFR-). Phenotyping classification into pro-inflammatory (CD206+) populations was performed. Analysis was conducted using FlowJo11 software (Version 11.1.0).

#### 2.10. Antibody panel and data acquisition

Cardiac and splenic cells in single suspension were stained with a 16-color myeloid-focused spectral flow cytometry panel: CD115/CSF-1R (BV421), CD11b (BUV615), CD11c (BV750), CCR2 (BUV661), CD45 (APC-Cy7), CD68/SR-D1 (BV711), CD86 (BV480), F4/80 (BUV563), Ly6G (RB780), Ly6G/Ly6C/Gr-1 (SB780), LYVE1 (AF488), Timd4 (PE), CD64 (PE-Cy5), MHC-II (eF506), CD161/NK1.1 (BV605), and Ghost Dye UV450 for viability. Data were acquired on a Cytek Aurora spectral cytometer.

#### 2.11. Data preprocessing

Spectral unmixing and compensation were performed in FlowJo v10.10.0. Events were sequentially gated for intact cells (FSC-A vs SSC-A), singlet discrimination (FSC-A vs FSC-H), viability (Ghost Dye UV450), and CD45+ selection (APC-Cy7) (Supplementary Figure 1A–B). For cardiac tissue, gating thresholds were adjusted to minimize inclusion of autofluorescent cardiomyocytes, confirmed by comparison with unstained controls. Compensated, scaled values for CD45+ events were exported as CSV files, yielding a combined dataset of 933,463 cells across 15 markers from 16 samples (4 groups × 4 replicates, paired heart and spleen per animal).

#### 2.12. Unsupervised clustering with CAFE

High-dimensional analysis was performed using CAFE (Cell Analyzer for Flow Experiments), an open-source Python application built on the Scanpy framework for GUI-based analysis of spectral flow cytometry data (Siam et al., Bioinformatics, 2025). Expression data were scaled (max value = 10) and reduced by PCA (10 components, auto SVD solver). Four batch correction methods (No Correction, ComBat, Harmony, BBKNN) were compared using Calinski-Harabasz, Davies-Bouldin, batch average silhouette width, and batch entropy metrics (Supplementary Figure 1D). Harmony was selected based on the highest clustering quality (Calinski-Harabasz = 87,712) and best tissue integration (batch ASW = −0.036), and was applied using cosine distance (20 maximum iterations). A shared nearest neighbor graph was constructed (n_neighbors = 30, cosine metric), and UMAP embedding was computed (min_dist = 0.1, spread = 1.0). Optimal clustering parameters were determined by systematic sweep across Leiden resolutions (0.1–1.0), neighbor counts, minimum distances, and distance metrics (Supplementary Figure 1C). Leiden clustering at resolution 0.5 (iGraph algorithm, 2 iterations) identified 24 distinct cell clusters, visualized by dot plots with hierarchical clustering dendrograms. Unsupervised clustering was used to characterize the overall immune cell landscape, while manual gating was employed to quantify predefined myeloid populations with sufficient statistical power.

#### 2.13. Manual gating analysis

In parallel, conventional biaxial gating in FlowJo quantified cardiac myeloid populations. CD45+ leukocytes were sequentially gated for non-myeloid markers, CD11b, and Ly6G (neutrophils). CD11b+Ly6G− cells were further assessed for CCR2, CD86, F4/80, and CD64. Populations were quantified as a percentage of CD45+ cells. Data are presented as mean ± SEM. [Specify statistical test]. *P < 0.05.

#### 2.14. Inflammatory assay in mouse bone marrow-derived macrophages

Mouse bone marrow-derived cells were extracted from the tibia and femur of both hind legs as previously described (24) and differentiated for 6-7 days in standard 1X DMEM containing 20% L929-conditioned media supplemented with 10% fetal bovine serum (FBS), 1% each of glutamax, pyruvate, and penicillin/streptomycin. Cells were stripped using the cell stripper (#25-056-Cl, Corning), counted, and plated into either 12-well or 24-well culture plates (#29443-952, Corning) to reach 90% confluency. Cells were allowed to settle overnight before being treated with PF8380 (10 nM and/or 500nM) for the next 24 hours. Cells were then stimulated with LPS (50 ng) for 6 hours before harvesting cells and supernatant for RT-PCR and ELISA, respectively. The total RNA was extracted using the standard TRIzol Reagent (#15-596-026, Invitrogen, USA) method, and RNA quality and concentration were assessed with a spectrophotometer (BioTek Synergy A1, Agilent Technology, USA). cDNA was synthesized using the iScript cDNA Synthesis kit (#1708890, BioRad, USA) using a thermal cycler (#MiniAmp Plus Thermal Cycler, Applied Biosystems), and multiple marker genes of inflammatory, anti-inflammatory, pro-fibrotic, and anti-fibrotic, including Rn18s as housekeeper genes, were amplified using PowerUp SYBR Green master mix (A25778, Thermo Fisher Scientific, USA).

#### 2.15. NFkB translocation assay in mouse bone marrow-derived cells

Mouse bone marrow-derived cells were extracted and plated in 12-well plate as stated above. Cells were allowed to adhere to the plate overnight before being exposed to PF8380 (500nM) for 24 hours. Next, cells were activated with lipopolysaccharide (LPS, 100 ng) and/or vehicle for 6 hours. Cells were then washed three times for 5 min each with cold 1X PBS, fixed with 4% PFA at room temperature for 15 min, and permeabilized with 0.3% Triton X100 for 10 min. Cells were washed again with 1X PBS to remove excess fixatives and permeabilization reagents before incubating with a 1:500 dilution of NF-κB p65 rabbit monoclonal antibody (D14E12, 8242s, Cell Signaling Technologies, USA) overnight at 4 °C. Following primary antibody incubation, cells were incubated with a 1:10,000 dilution of goat anti-rabbit secondary antibody (#ab150081, Alexa Fluor 488) against the primary for 1 hour at room temperature. Coverslips containing cells were washed in the wells, transferred to the slides, and imaged using a Keyence fluorescent microscope (Keyence, BZ-X810, Japan).

#### 2.16. Statistical analysis

All statistical analyses were performed on either RStudio 2026.01.0 (Posit Software, PBC) or GraphPad Prism 10 for macOS (Version 10.6.1, GraphPad Software, LLC), and data are presented as mean±SEM. Normality check on the data was performed using Shapiro-Wilk test to confirm the normal distribution and parametric test was applied on log-transformed data if not normally distributed. The human lipidomic data was read into a *Summarized Experiment* object and matched with the clinical data on ID through *dplyr*. The lipid shorthands were canonicalized using the *RefMet* database followed by *rgoslin* for maximum coverage. The lipidomics data was then filtered for 70% availability and normalized by percent log-base-10 fold change using *LipidSigR*. Differential analysis, as represented on the volcano and lollipop plots, was then conducted. A correlative analysis heatmap was then produced using the same data processing workflow. All plots were rendered using *ggplot2* and *ggrepel*. For mouse data, one-way ANOVA was performed between groups to compare the means between groups (diet+drug) as a main factor between groups, with Tukey’s post hoc test applied. P<0.05 was considered statistically significant.

## Results

### Heart Failure with Preserved Ejection Fraction (HFpEF) patients have higher circulating levels of pro-inflammatory cytokines and lipid signature

Half of all heart failure cases are characterized by sterile low levels of systemic inflammation (4), and inflammatory markers are usually associated with left ventricular diastolic dysfunction (6, 7). We conducted an assessment on total plasma levels of LPA level including Luminex assay, a multiplex assay to assess the systemic inflammatory landscape in HFpEF. Patients with HFpEF had significantly higher plasma levels of LPA **(Fig. 1B)** including all nine inflammatory mediator proteins tested in human plasma samples, namely interleukin-6 (IL-6), Syndecan-1, endonucleotide pyrophoaphatase/phosphodiesterase 2 (ENPP2), fibroblast growth factor-23 (FGF-23), growth differentiation factor (GDF-15), Syndecan-4, Galectin-3, c-reactive protein (CRP), tumor necrosis factor-alpha (TNF-α), were significantly upregulated in HFpEF patients **(Fig. 1C-1K),** a finding that is consistent with the previous reports (4, 5). Previous studies including ours have shown that bioactive lipids such as lysophosphatidic acid (LPA) and its associated G-protein coupled receptors (GPCRs, LPA1-6) do not just govern systemic sterile levels of inflammation (25), but also modulates the myeloid cells response to injury during myocardial infarction (14–16) and helps in development and progression of cardiac (13) and pulmonary fibrosis (18). To understand the potential molecular role of bioactive lipids, including LPA-specific lipids, in HFpEF development, human plasma samples were subjected to mass spectrometry for both targeted (LPA-specific) and untargeted lipidomic analysis. Differential lipid abundance analysis visualized by a volcano plot revealed a distinct pattern of lipid remodeling that is associated with maintaining the structural integrity of the cell membrane, cell signaling and a measure to prevent metabolically driven stress-related inflammation driven by metabolic shift during HFpEF. Our findings indicate that lysophosphatidylethanolamine 18:2 (LPE O-18:2), including ether-linked LPE O-16:1, phosphatidylserine, and phosphatidylethanolamine, are upregulated in HFpEF compared to control (|-log10 fold change| ≥0.5-1.5, FDR<0.05) **(Fig. 1L)**. These HFpEF patients also have a significantly higher levels of phosphatidylcholine-36:4;O, an ether-linked lipid **(Fig. 1M)** and significantly downregulated sphingomyelin-39:2; O3 (SM 39:2; O3), and phosphatidylserine-36:0 (PS 36:0) **(Fig. 1M)**. Majority of these bioactive lipids were upregulated in HFpEF patients with hypertension, type 2 diabetes, and were under hyperlipidemic drug **(Fig. 1N)** – the clinical features that defines HFpEF. Given that more than half of HFpEF patients are female (1), we stratified the lipidomic findings by gender to assess sexual dimorphism in the lipid signature defining HFpEF. Interestingly, both male and female shares significant increase in similar lipid profile (such as sterol lipid (ST 27:1; O;S), LPC, PA, ceramide (Cer), and acylcarnitine (CAR)) that are elevated with advancing age, and metabolic phenotype such as hypertension, type 2 diabetes and among the patients those who were on lipid lowering drug **(Fig. 2A** and **B)**. Moreover, some of the major bioactive lipids, such as phosphatidylserine 38:4 (PS38:4) and LPE 16:1, were even higher in females compared to males, even though they share a similar level of PC36:4; O and LPE O-18:2 **(Fig. 2C, D, E,** and **F)**. We further examined our specific lipids using the lollipop plot, revealing an upregulated PE subclass such as PE 36:5 and PE 40:8, previously not observed in the volcano plot **(Fig. 2F),** suggesting PE is a shared upregulated lipid in HFpEF in both male and female patients. This is interesting considering the role of PE in upregulation of autophagy via lipidation of LC3I to LC3II – a process commonly known as the cellular self-cleansing mechanism (26).

**Figure 1.**
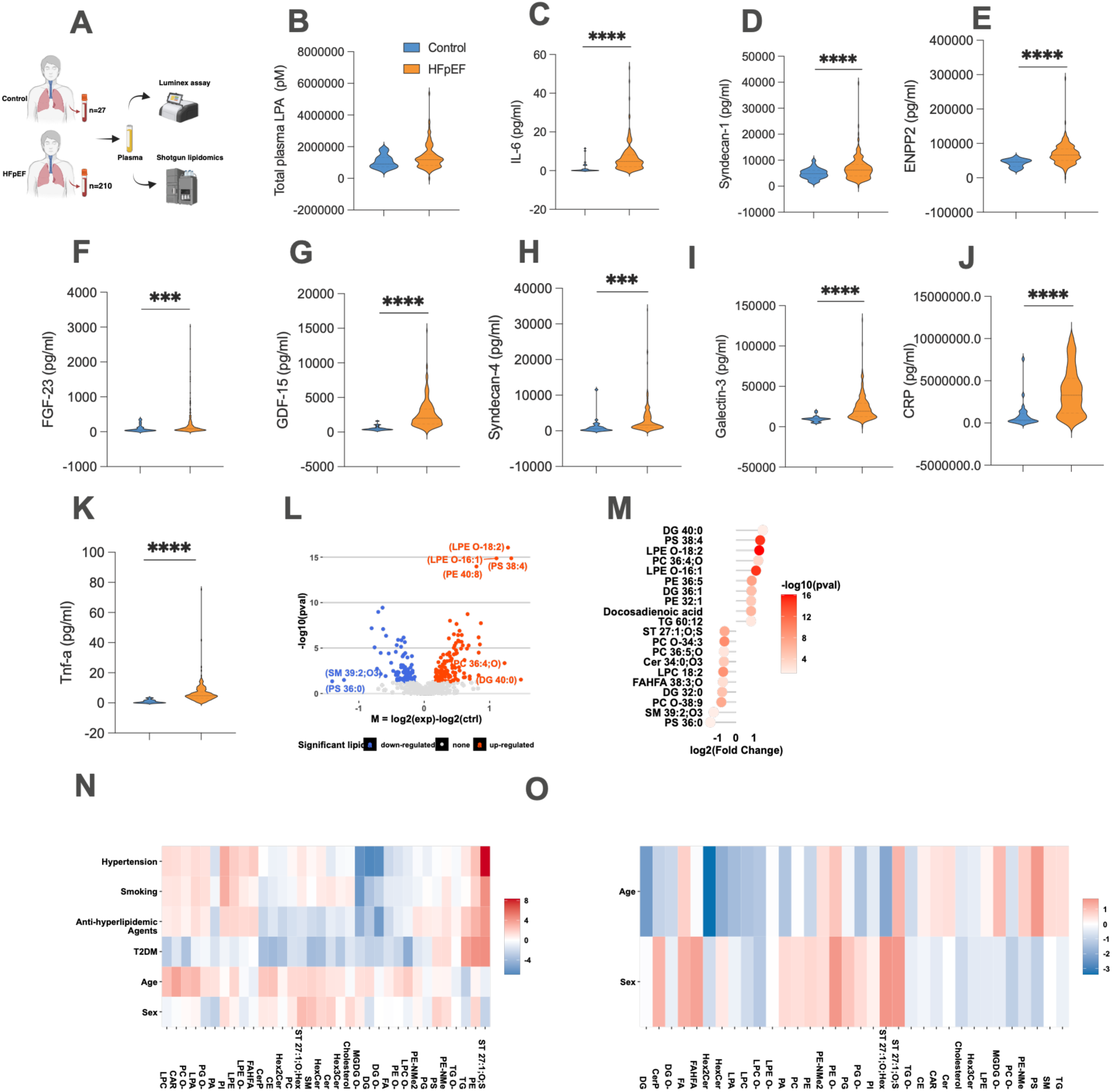
Pro-inflammatory cytokines and lipid signatures are higher in HFpEF compared to controls. **A.** Experimental design of human study. **B.** Total plasma levels of LPA in human HFpEF (n=140) and Controls (n=27) in picomolar **C- K**. Plasma levels of inflammatory proteins in HFpFF (n=210) vs Controls (n=27). **L.** Differential lipidomic analysis visualized with volcano plot in HFpEF (n=210) vs Controls (n=27). **M.** Differential analysis of the untargeted lipidome represented in volcano and lollipop plots with very highly and lowly expressed lipid classes. **N.** Heatmap representation of the correlation analysis of the differentially expressed lipids in relation to the clinical and anthropometric data in HFpEF (n-210). **O.** Heatmap representation of the correlation analysis of the differentially expressed lipids in relation to the age and sex in control group (n=27). **HFpEF:** Heart failure with preserved ejection fraction. **IL-6:** Interleukin-6, **ENPP2:** Ectonucleotide pyrophosphatase/phosphodiesterase 2, **FGF-23:** Fibroblast growth factor-23, **GDF-15:** Growth differentiation factor-15, **CRP:** C-reactive protein, **Tnf-a:** Tumor necrosis factor-alpha. **pM:** picomolar *P*<0.05, *\*\*P*<0.01, *\*\*\*P*<0.001, *\*\*\*\*P*<0.0001. **RED:** significantly high expressed lipids. **Blue:** significantly low expressed lipids.

**Figure 2.**
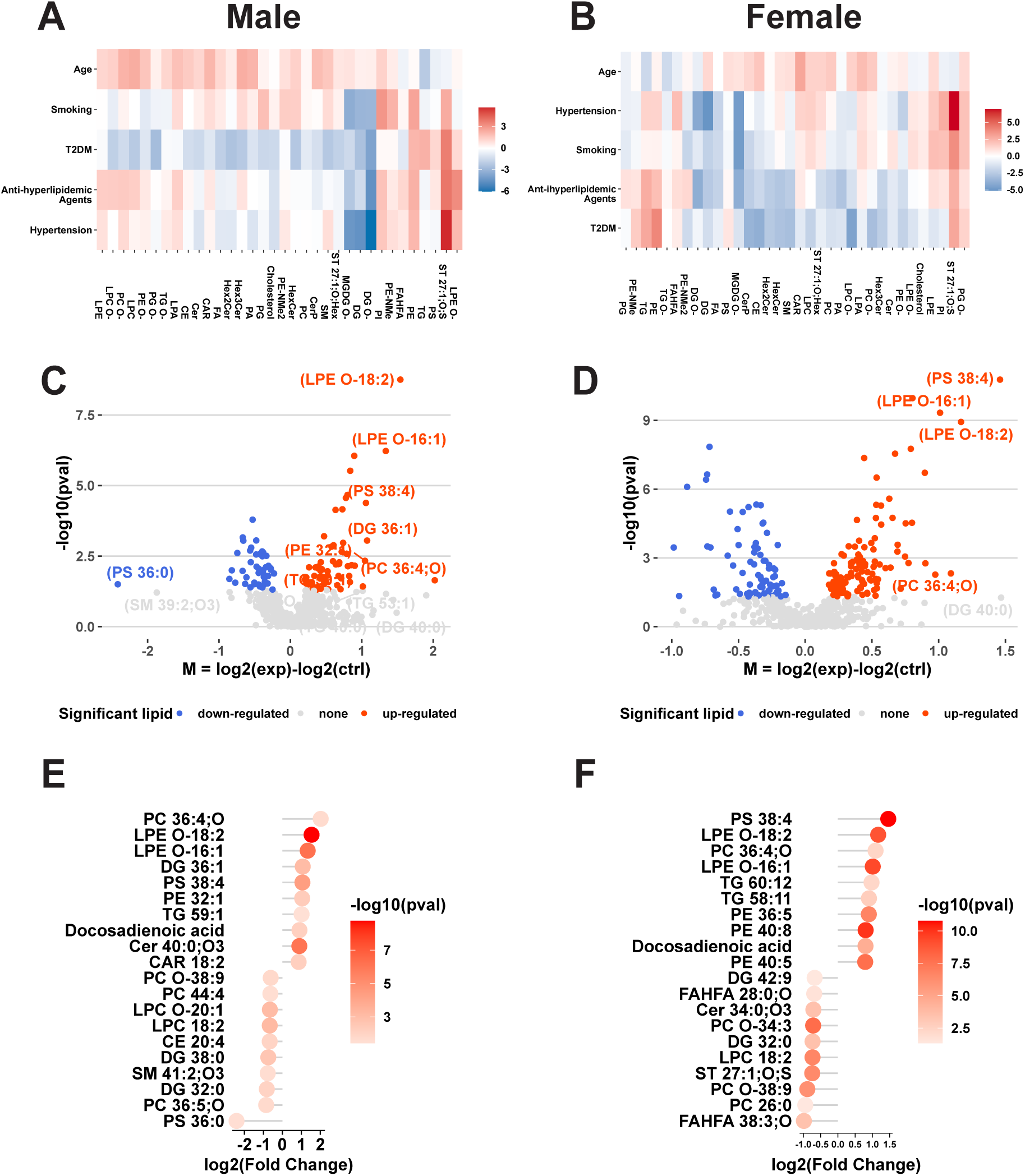
Sex-mediated lipidomic dimorphism in human HFpEF compared to controls. **A.** Heatmap representation of correlation analysis of anthropometric and clinical data with lipids in male HFpEF. **B.** Heatmap representation of correlation analysis of anthropometric and clinical data with lipids in female HFpEF. **C.** Volcano plot of log-transformed data of differentially expressed plasma lipids in male HFpEF. **D.** Volcano plot of log-transformed data of differentially expressed plasma lipids in female HFpEF. **E.** Lollipop plot of log2 (fold change) of very high and low differentially expressed lipid classes in male HFpEF. **F.** Lollipop plot of log2 (fold change) very high and low differentially expressed lipids classes in female HFpEF. **RED:** significantly high expressed lipids. **Blue:** significantly low expressed lipids. **Dark orange:** very high expressed lipid class. **Gray:** very low expressed lipid class.

### PF8380 administration for 10-weeks reduces metabolic phenotypes in HFpEF mouse model

Increased de novo synthesis and/or accumulation of pro-inflammatory bioactive lipids, such as ceramide, are associated with adverse cardiac remodeling (27). Additionally, studies, including ours, have shown that the lysophosphatidic acid (LPA)-autotaxin axis promotes fibrosis in major organs such as the heart (16), lung (11), and kidney (28). Interestingly, our untargeted lipidomic analysis revealed significant upregulation of several major inflammatory lipid mediators in both male and female HFpEF patients, and, more importantly, LPE, the second major substrate after lysophosphatidylcholine for autotaxin to produce lysophosphatidic acid (LPA). Based on these observations, we hypothesized that the LPA-autotaxin axis may be a major contributor to the pathophysiology of HFpEF. To confirm this, we used PF8380 against autotaxin for 10 weeks in an established mouse HFpEF model confirmed through a series of experiments at week 5 **(Fig. 3A-F)**. HFpEF mice developed through a modified two-hit model had increased body weight **(Fig. 3A** and **B)**, elevated systolic and diastolic blood pressure **(Fig. 3B** and **C),** and significantly reduced glucose tolerance **(Fig. 3E** and **F)**. As expected, ten weeks of PF8380 administration reduced systolic and diastolic blood pressure and improved glucose tolerance. Although total body weight at week-15 was reduced in the PF8380 group (**Fig. 3G-L**), we did not find a significant difference in body fat mass and/or lean mass, including their total percentage composition, between the PF8380 vs vehicle group when assessed by EchoMRI **(Supplementary Figure 2, A-J)**. Further, exercise exhaustion test revealed a significant improvement in the total running time and the distance covered by the PF8380 group mice at week-15 compared to the vehicle group (**data not shown**).

**Figure 3.**
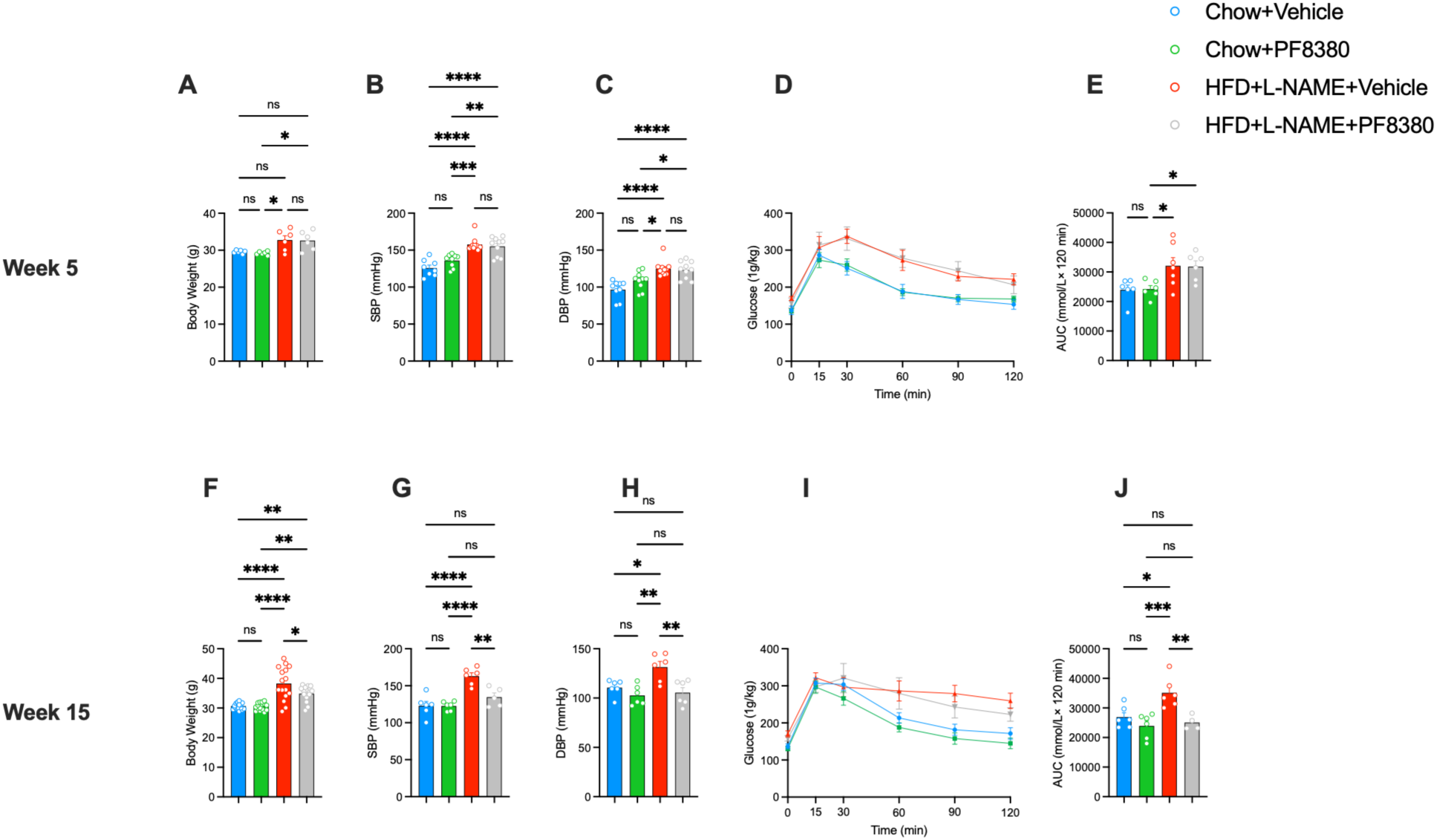
PF8380-mediated autotaxin inhibition reduces metabolic parameters in C57BL/6J mouse models with the HFpEF phenotype. **(A)** Weekly body weight in grams at week 5 in C57BL/6J mice (6/group). **(B)** Systolic blood pressure at week 5 (8/group). **(C)** Diastolic blood pressure at week 5 (8/group). **(D, E)** Intraperitoneal blood glucose tolerance test (IPGTT, 1g/kg body weight) (6/group). **(F)** Weekly body weight at week 15 (12/group). **(G)** Systolic blood pressure at week 15 (6/group). **(H)** Diastolic blood pressure at week 15 (6/group). **(I, J)** Intraperitoneal blood glucose tolerance test at week 15 (6/group). One-way ANOVA with Tukey’s post-hoc correction. *\*P*<0.05, *\*\*P*<0.01, *\*\*\*P*<0.001, and *\*\*\*\*P*<0.0001. **SBP:** Systolic blood pressure, **DBP:** diastolic blood pressure, **AUC:** area under the curve, mm: millimole.

### PF8380 administration against autotaxin prevents adverse cardiac remodeling in a mouse model of HFpEF

Existing evidence suggests that mice exposed to a high-fat diet in combination with L-NAME develop common functional and structural features of human HFpEF observed in clinical settings. Based on these observations, we exposed the mice to HFD+L-NAME for week-5 to develop the HFpEF phenotype. Mice were confirmed to have an HFpEF phenotype at week 5 through a series of experiments, including echocardiography, an intraperitoneal blood glucose tolerance test, and tail-cuff blood pressure measurements. Mice then received daily oral gavage of PF8380 against autotaxin (3 mg/kg body weight) for another 10-weeks. Mice receiving HFD+L-NAME in combination had an increased plasma levels of LPA at week-5 before administration of PF8380 against autotaxin **(Fig. 4B)**. Echocardiographic examination revealed a decrease in E/A ratio, isovolumic relaxation time in the PF8380 group, and left ventricle anterior wall diametercompared to the vehicle group **(Fig. 4E, 4G,** and **4H**, respectively) that was otherwise increased with the exposure to HFD+L-NAME dietary regimen. Cardiac fibrosis is a hallmark feature of HFpEF in both humans (29) and mice (30). Mice’s hearts were then harvested to assess structural and molecular changes observed at week-15 in all four groups. Masson’s trichrome staining revealed a significant deposition of collagen fiber in the mouse heart **(Fig. 4I** and **J)** that was mitigated by 10 weeks of PF8380 administration **(Fig. 4I** and **4J)**. Further image analysis revealed a significant decrease in both cardiomyocyte size **(Fig. 4K** and **4L)** and perimeter **(Fig. 4M)** in the mouse heart following PF8380 treatment. PF8380 increased the levels of *Anp* gene expression in mouse heart tissue upon further analysis using RT-qPCR method **(Fig. 4N)**.

**Figure 4.**
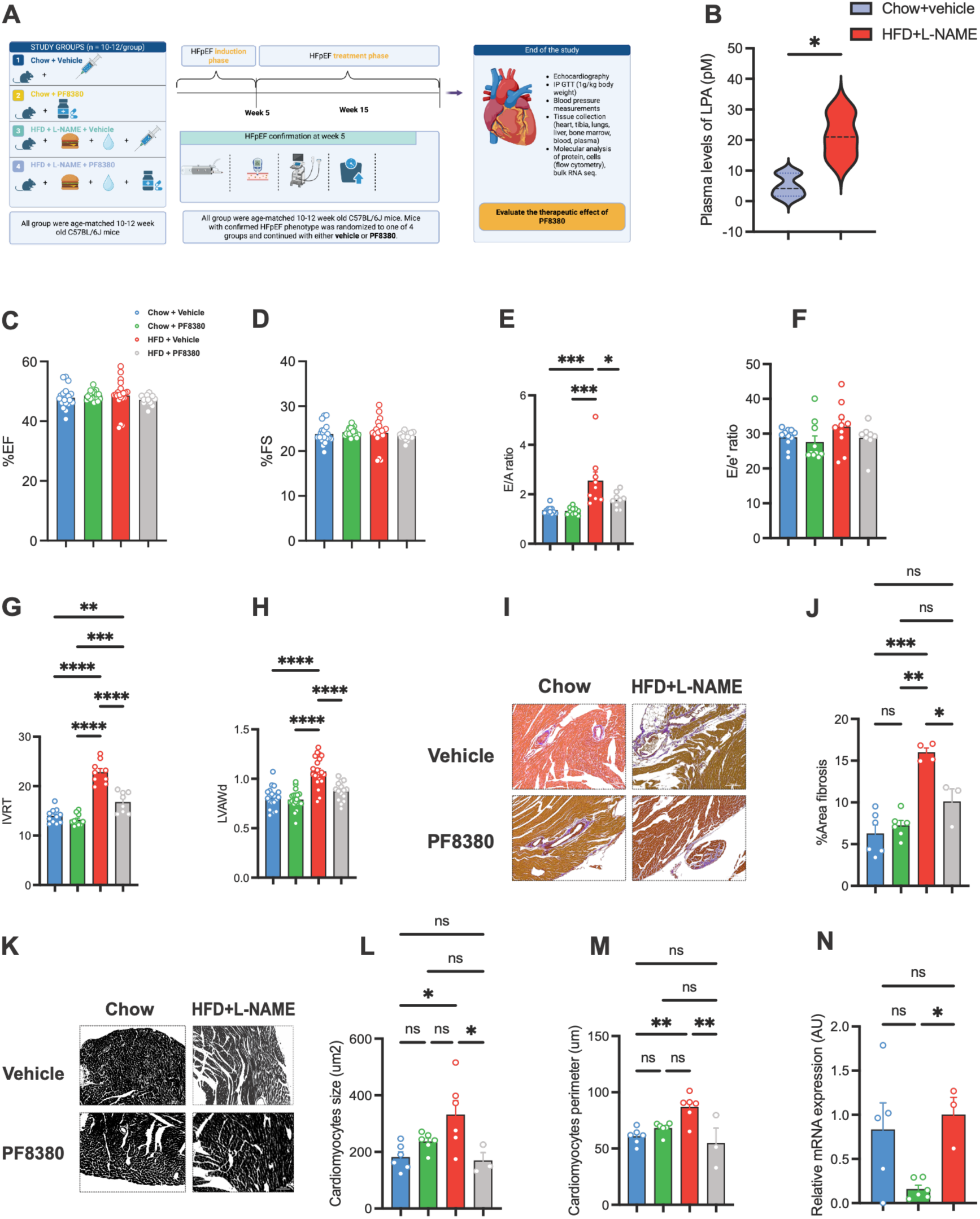
Mouse cardiac structure and function were assessed at week 15. **(A)** An animal experimental design was maintained at different dietary regimens for 15 weeks. **(B)** Plasma levels of LPA in Chow (n=6/group) and HFD+L-NAME mice (n=3/group). **(C)** Percentage ejection fraction at week-15 (n=12/group). **(D)** Percentage fractional shortening at week-15 (n=12/group). **(E)** E/A ratio of the mouse heart at week-15 (n=10/group). **(F)** E/e’ ratio at week-15 (n=10/group). **(G)** Isovolumic relaxation time of the mouse heart at week-15 (n=12/group). **(H)** Left ventricle anterior wall diameter at week-15 (n=12/group). **(I** and **J)** Representative images of Masson’s trichrome stain of the mouse heart at week-15 (n=3-6/group), and quantification as percentage fibrosis of the mouse heart at week-15 (n=3-6/group). **(K)** Representative image of cardiomyocyte size at 20X magnification using MTC-stained images at week-15 (n=3-6/group). **(L** and **M)** Quantification of cardiomyocytes size in μm^2^ and perimeter in μm at week-15 (n=3-6/group). **(N)** Relative mRNA *Anp* expression in cardiac tissue (3-4/group). One-way ANOVA with Tukey’s post-hoc test. *P<0.05, **P<0.01, ***P<0.001, and ****P<0.0001.

### Unsupervised clustering reveals a heterogeneous immune cell landscape in cardiac and splenic tissues of HFpEF mice

To characterize the immune cell composition of cardiac and splenic tissues in the two-hit HFpEF model, we performed unsupervised analysis of 16-color spectral flow cytometry data using CAFE (Siam et al., Bioinformatics, 2025). Following quality control gating, Harmony-based batch correction, and optimization of clustering parameters (**Supplementary Figure 1**), Leiden clustering identified 24 distinct cell populations across 933,463 CD45+ events from heart and spleen **(Fig. 5A)**. Hierarchical clustering of marker expression profiles revealed that cardiac and splenic tissues harbored distinct immune cell compositions with partially overlapping cluster identities (**Fig. 5A**). UMAP visualization confirmed clear tissue-dependent segregation of clusters, with several populations enriched preferentially in the heart (clusters 0, 1, 2) or the spleen (clusters 5, 8, 9, 11, 12, 15), while others were shared across both compartments (**Fig. 5B–C**). Dot plot analysis of marker expression revealed tissue-specific patterns: cardiac-enriched clusters expressed high levels of F4/80, Timd4, and LYVE1, consistent with tissue-resident macrophage identity, while splenic clusters displayed greater heterogeneity in myeloid and non-myeloid marker expression, including CD11b, CCR2, CD86, and MHCII (**Fig. 5A**). UMAP overlay by treatment group indicated representation of all four conditions (Chow+Vehicle, Chow+PF8380, HFD+L-NAME+Vehicle, HFD+L-NAME+PF8380) across the identified clusters **(Fig. 5C)**.

**Figure 5.**
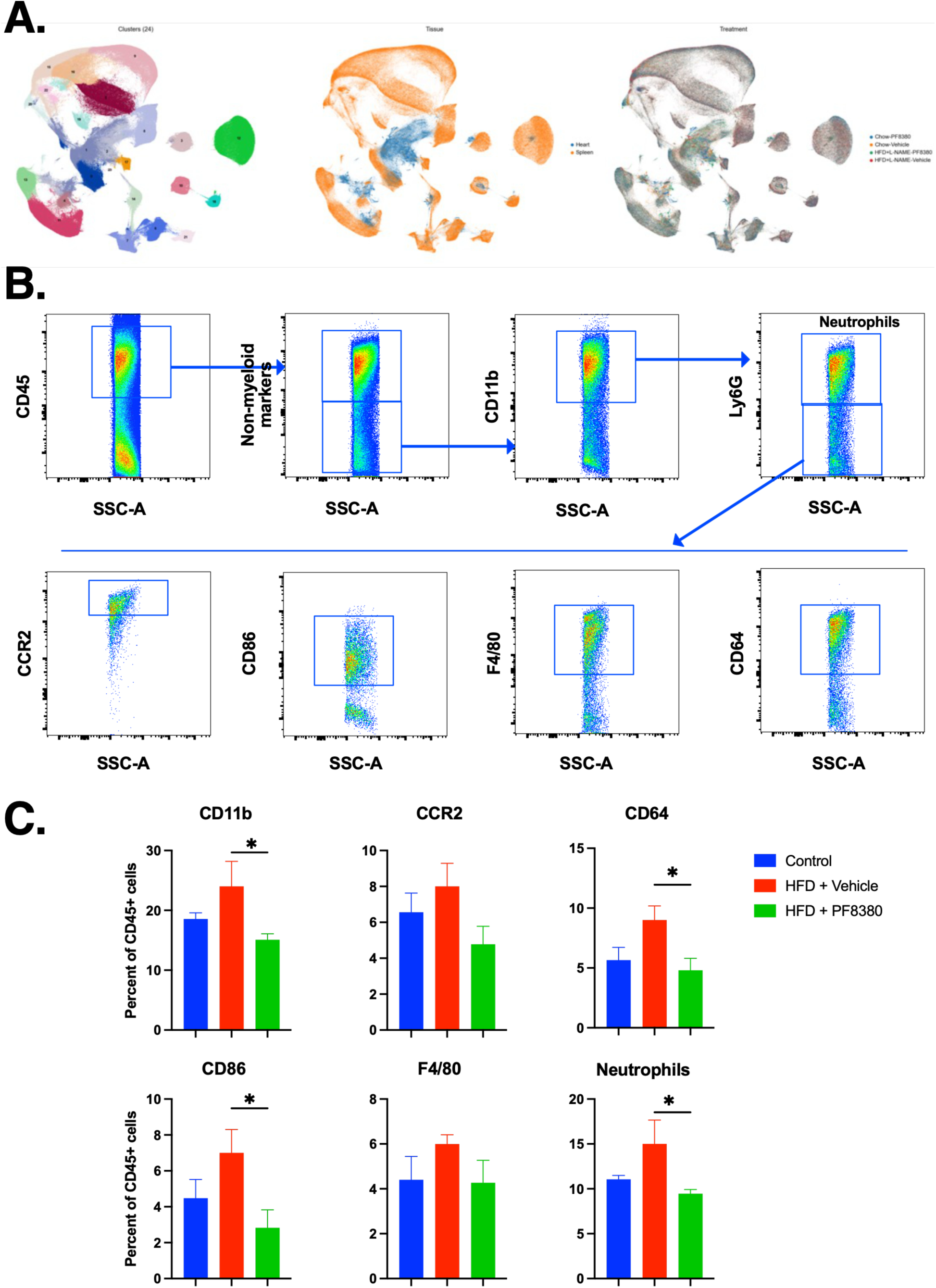
Unsupervised and manual gating analyses of cardiac and splenic immune cell populations in a two-hit HFpEF model treated with the autotaxin inhibitor PF8380. **(A)** Dot plots displaying marker expression profiles across Leiden clusters identified by unsupervised analysis using CAFE (Cell Analyzer for Flow Experiments). Dot size represents the fraction of cells expressing each marker within a cluster (%), and color intensity indicates mean expression level (scaled 0–1). Hierarchical clustering dendrograms group clusters by expression similarity. Data are shown for combined heart and spleen (left), spleen only (center), and heart only (right). Markers analyzed include CD45, LYVE1, F4/80, CD11b, CCR2, CD115, CD86, CD161, CD11c, CD68, Timd4, CD64, Ly6G, Ly6G/Ly6C, and MHCII. **(A)** UMAP (Uniform Manifold Approximation and Projection) visualization of 24 Leiden clusters (numbered 0–23) in heart (blue) and spleen (orange) tissues. Each cluster is represented by a distinct color. UMAP projections of combined heart and spleen data colored by cluster identity (24 clusters; left), issue of origin (heart, blue; spleen, orange; center), and treatment group (Chow+PF8380, Chow+Vehicle, HFD+L-NAME+PF8380, HFD+L-NAME+Vehicle; right). **(B)** Representative manual gating strategy for identification of cardiac myeloid cell populations. CD45+ leukocytes were gated from live, single-cell events and sequentially classified based on non-myeloid lineage markers and CD11b expression. CD11b+ cells were assessed for Ly6G to identify neutrophils. CD11b+Ly6G− cells were further analyzed for CCR2, CD86, F4/80, and CD64 expression to characterize monocyte/macrophage subsets. **(C)** Quantification of myeloid cell populations expressed as percent of CD45+ cells in cardiac tissue across three groups: Control (blue), HFD + Vehicle (red), and HFD + PF8380 (green). For panels A–C, spectral flow cytometry data were acquired using a 16-color panel Supplementary Table 1). Compensated, scaled data were exported from FlowJo v10.10.0 following gating on live, single CD45+ cells. Unsupervised analysis was performed using CAFE (Siam et al., Bioinformatics, 2025). Data were scaled (max value = 10), reduced by PCA (10 components, auto SVD solver), batch-corrected using Harmony (cosine metric, 20 maximum iterations), and projected by UMAP (n_neighbors = 30, min_dist = 0.1, spread = 1.0). Community detection was performed by Leiden clustering (resolution = 0.5, iGraph algorithm, 2 iterations), yielding 24 distinct clusters. Data presented as mean ± SEM. *P < 0.05; One-way ANOVA with Tukey’s post hoc test. n = 4/group.

### PF8380-mediated autotaxin inhibition attenuates cardiac myeloid cell accumulation in HFpEF

To quantify treatment effects on defined myeloid populations, we manually gated cardiac CD45+ leukocytes using established monocyte, macrophage, and neutrophil markers (**Fig. 5B**). HFD + Vehicle mice exhibited increased CD11b+ myeloid cells compared to controls, and this increase was significantly attenuated by PF8380 treatment (**Fig. 5B**). Among myeloid subsets, CD64+ macrophages and CD86+ activated myeloid cells were significantly elevated in HFD + Vehicle hearts relative to HFD + PF8380-treated animals (**Fig. 5B and 5C**). Cardiac neutrophil frequency (Ly6G+) was also significantly higher in HFD + Vehicle compared to HFD + PF8380-treated mice (**Fig. 5B**). CCR2+ monocytes and F4/80+ macrophages showed a similar directional trend, increased in HFD + Vehicle and reduced by PF8380, though these differences did not reach statistical significance (**Fig. 5B** and **5C**). Collectively, these data indicate that autotaxin inhibition reduces cardiac myeloid cell infiltration in the two-hit HFpEF model, with significant effects on total myeloid cells, macrophages, activated myeloid cells, and neutrophils.

### PF8380 attenuates LPS and/or LPA-mediated inflammation in mouse macrophage via NFkB pathway

We and others have previously shown that pharmacological and/or genetic inhibition of ATX/LPA signaling axis attenuates inflammatory response after acute myocardial infarction (16, 31). Based on these observations, we hypothesize that PF8380 may have broader anti-inflammatory potential, given its ability to reduce LPA production via autotaxin-mediated hydrolysis of lysophosphatidylcholine. To mimic the inflammatory milieu observed in HFpEF conditions in an *invitro* model, mouse bone marrow-derived cells were subjected to PF8380 for 24 hours before being stimulated with either LPS (50 ng and/or 100ng, #00-4976, Invitrogen) or LPA (10 nM, #A85128, Avanti Research, USA) for 6 hours. As expected, PF8380 significantly prevented the translocation of NFkB into the nucleus in the cells as assessed by the ratio of nuclear to cytoplasmic NFkB **(Fig. 6B)** and reduced *TNF-α* levels **(Fig. 6C)** and increased both IL-10 and IL-6 expression **(Fig. 6D)** in the macrophage supernatant when compared to the LPS group.

**Figure 6.**
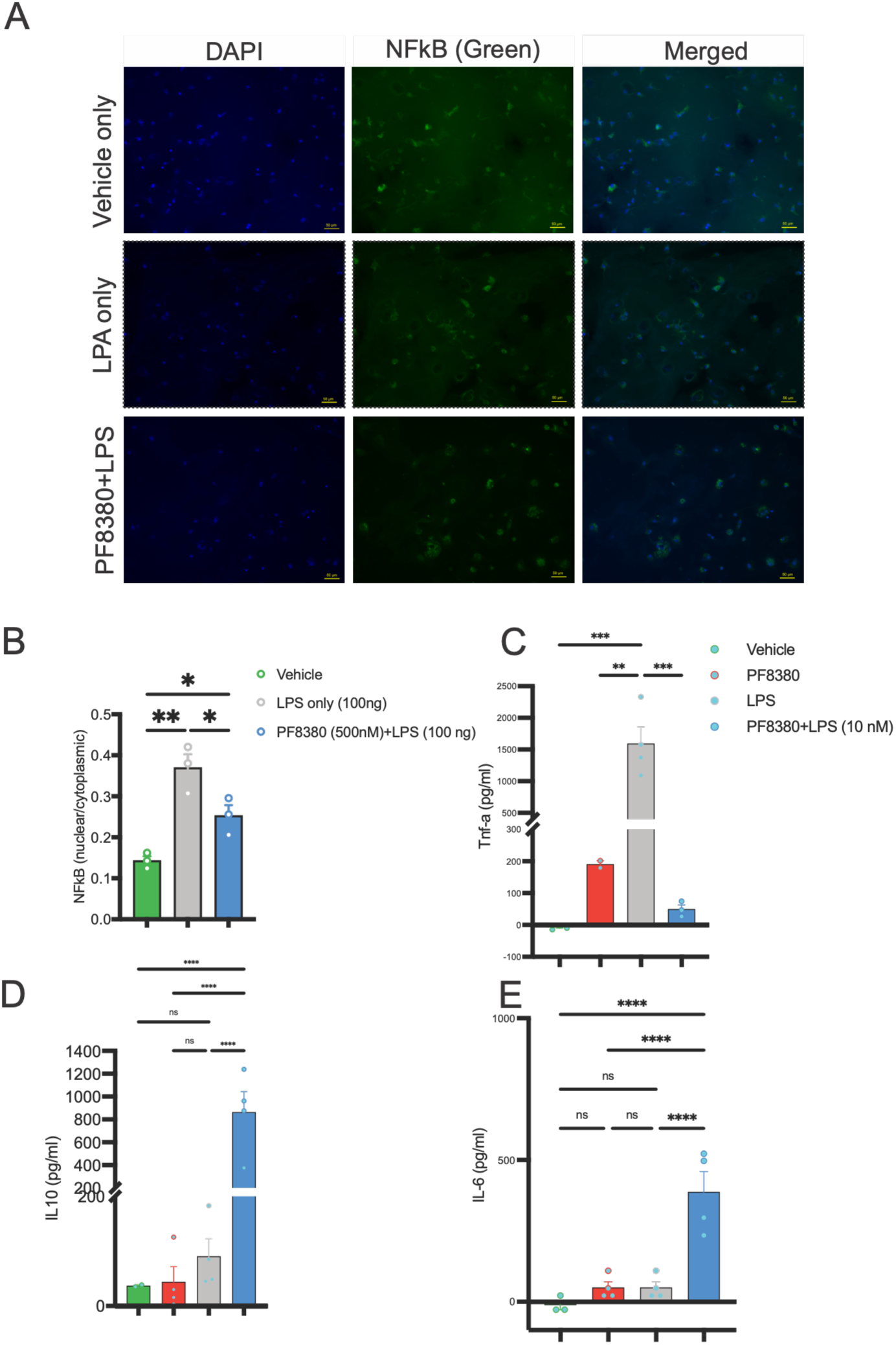
PF8380-driven NFkB-mediated anti-inflammatory assay in mouse primary bone marrow-derived cells. **(A)**. Staining of nucleus (DAPI, blue, left column), NFkB (green, middle column) in mouse primary bone marrow-derived cells treated with vehicle, LPS only (100 ng), and PF8380 (500 nM) with LPS (100 ng) activation, scale bar 50 μm. **(B)** Ratio of nuclear/cytoplasmic nuclear factor kappa-b (NFkB) in mouse BMDM (n=3/group). **(C)** Tnf-α expression in mouse BMDM supernatant in picograms per ml (n=2-4/group). **(D)** IL-10 expression in mouse macrophage in picograms per ml in mouse BMDM supernatant (n=2-4/group). **(E)** IL-6 expression in mouse macrophage in picograms per ml in mouse BMDM (n=3-4/group). One-way ANOVA with Tukey’s post hoc test. *P<0.05, **P<0.01, ***P<0.001, and ****P<0.0001.

## Discussion

HFpEF remains a leading cause of morbidity and mortality worldwide, and the persistent lack of disease-modifying therapies underscores the need to identify new mechanistic targets. In this study, we tested whether the autotaxin-LPA axis is engaged in heart failure with preserved ejection fraction (HFpEF) and whether pharmacologic inhibition of this axis is therapeutic. Three principal findings emerged. First, patients with HFpEF exhibited a coordinated elevation of circulating inflammatory lipid mediators and a remodeled plasma lipidome enriched for autotaxin substrates, with autotaxin (ENPP2) itself among the upregulated markers. Second, autotaxin inhibition with PF8380 in a two-hit (high-fat diet plus L-NAME) murine model of HFpEF improved diastolic function and enhanced exercise capacity. Third, these functional benefits were accompanied by reduced cardiac fibrosis and cardiomyocyte hypertrophy, and diminished myeloid cell accumulation in the myocardium. Together, these data identify the autotaxin-LPA axis as an inflammatory driver of HFpEF and establish inhibition of this axis as a therapeutic strategy.

The prevailing model holds that the metabolic comorbidities of HFpEF, namely obesity, hypertension, and diabetes, converge on a state of systemic inflammation that produces coronary microvascular dysfunction and myocardial stiffening (Sanders-van Wijk et al., 2020; Verma et al., 2024). However, large clinical trials such as phase II ENDEAVOR targeting systemic inflammation via myeloperoxidase inhibition failed to improve overall cardiac health and therefore functional capacity of the patients (32). Autotaxin is a phospholipase D enzyme, secreted primarily by the adipose tissue (33) that converts lysophosphatidylcholine to lysophosphatidic acid (LPC), one of the major components that drives systemic level of inflammation and aids in fibrosis in skeletal muscle (34), kidney (12), and pulmonary fibrosis (35). Central obesity is a major comorbidity of HFpEF (36). Based on this consensus, we hypothesized that obesity-mediated autotaxin release from adipose tissue converts circulating lysophosphatidylcholine (LPC) to lysophosphatidic acid (LPA), contributing to systemic inflammation. In consensus with the previous studies, HFpEF patients in our study showed significantly higher levels of inflammatory markers in plasma samples (37–39). Fatty acids are the major source of energy for the heart (40); however, altered plasma lipidome and abnormal lipid metabolism are frequently associated with human HFpEF (41, 42). Although there is no reported lipidomic data on human HFpEF, female HFpEF patients usually have clinically diagnosed higher diastolic dysfunction compared to male (43), suggesting existence of sexual dimorphism in human HFpEF (44). Our findings indicates that HFpEF patients have significantly higher level of systemic inflammation with female HFpEF harboring even more pronounced pro-inflammatory and membrane-destabilizing lipid signature compared to male. Both sexes show an elevation of LPE/LPC and ether-lined species (LPE-O, PC-O) suggesting phospholipase activation, mitochondrial stress and altered membrane turnover – a mechanistic hallmark of HFpEF pathogenesis. Female HFpEF participants reflect higher lipid remodeling amplified by comorbidities such as hypertension and type 2 diabetes mellitus (T2DM) and very strong upregulation of PS38:4, LPE O-16:1, LPE O-18:2, PC 36:4;0. Female HFpEF participants have much larger number of significant lipids than male and points to same pathways activation as in male but with far stronger activation – this is consistent with the clinical HFpEF biology.

Human plasma lipidomic studies in HFpEF remain limited; our findings add extend altered lipidome knowledge on HFpEF to the previously reported data on HFpEF and the lipid-enriched metabolomic signature observed in patients with HFpEF and metabolic liver disease (41, 42). Both male and female HFpEF participants showed a concordant increase in LPE O-18:2, LPE O-16:1, PS 38:4, and PC 36:4;O, indicating that phospholipid remodeling is a common feature of HFpEF independent of sex. However, female HFpEF participants had a much larger number of statistically significant lipid alterations, with PS 38:4 and the two ether-linked LPE species representing some of the most strongly increased lipids, together with several polyunsaturated PE and TG species. Male HFpEF participants showed activation of similar pathways, although the overall signature was less extensive and included elevations in DG 36:1, PE 32:1, and TG 59:1. In consensus with population lipidomic studies showing that lysophospholipid and ether-phospholipid metabolism are strongly influenced by sex and age (45), our results suggest that female HFpEF patients have a higher phospholipid remodeling compared to male – this may contribute to the more pronounced diastolic dysfunction commonly observed in female HFpEF patients (43). The simultaneous remodeling of LPE, PC, PE, PS, DG, and TG species may reflect increased phospholipase-dependent membrane turnover, altered lipid reacylation, extracellular-vesicle or lipoprotein trafficking, and abnormal mitochondrial or peroxisomal lipid handling. However, the lipid changes were not uniform across individual classes; particularly, several LPC and ether-PC species were reduced rather than increased, especially in male HFpEF. Therefore, these results point toward broad and sex-dependent lipid remodeling rather than generalized accumulation of lysophospholipids. The marked changes in lysophospholipid point towards altered autotaxin-related lipid metabolism; therefore, we further analyzed the autotaxin-LPA axis by investigating the LPA-targeted lipidome.

Our targeted lipidomic findings provide direct evidence that circulating LPA metabolism is altered in human HFpEF. LPA 20:0 was similarly increased by approximately 3.5-fold in both male and female HFpEF participants. Female HFpEF participants demonstrated a broader LPA signature characterized by increases in LPA 16:1, LPA 20:0, LPA 22:0, LPA 22:2, and LPA 22:6, whereas LPA 18:2 was significantly reduced. This amplified signature likely reflects the greater comorbidity burden, notably hypertension and type 2 diabetes mellitus (T2DM), carried by women with HFpEF, which intensifies lipid remodeling and systemic inflammation. This is consistent with the clinical heart failure biology, where women have higher alterations in atherogenic lipids and systemic levels of inflammatory markers during type 2 diabetes and have a higher risk of coronary heart disease (46)Male HFpEF participants showed a more restricted pattern consisting primarily of increased LPA 20:0 and LPA 22:4. These results indicate that HFpEF does not produce a uniform elevation of the total LPA pool but instead alters its molecular-species composition, with preferential enrichment of selected saturated and very-long-chain LPA species. In combination with the increase in circulating LPE species identified in our untargeted lipidomic analysis, these findings support increased lysophospholipid remodeling in HFpEF and are compatible with activation of the autotaxin–LPA axis. However, because circulating autotaxin protein, enzymatic activity, and species-matched LPA/LPC ratios were not measured, further study is required to establish autotaxin as the direct source of the altered LPA profile. Future studies integrating autotaxin activity with absolute LPA and LPC concentrations will be necessary to determine whether this pathway contributes to systemic inflammation, vascular dysfunction, and fibrosis in HFpEF. Our current findings position the autotaxin-LPA axis as a specific, druggable node within this comorbidity-inflammation paradigm. The convergence of the human and murine data is notable given the fact that the same lipid-generating enzyme that was elevated in patient plasma, when inhibited in a mouse model of HFpEF, either reversed or mitigated major metabolic parameters that define and determine HFpEF pathophysiological conditions. Although the transability potential of mouse HFpEF findings in human has inherently been challenging across multiple disease models in past, the potential benefits of PF8380-mediated ATX inhibition against HFpEF in mouse model is assuring, considering the existence of clinical heterogeneity in human HFpEF (47). This concordance across species strengthens the inference that autotaxin-derived LPA is not merely a correlate of HFpEF but one of the major contributors to its pathogenesis (48).

In this study, our spectral flow cytometry showed that autotaxin inhibition attenuated the accumulation of total myeloid cells, macrophages, activated (CD86+) myeloid cells, and neutrophils in the HFpEF myocardium. In bone marrow-derived macrophages, PF8380 reduced LPS-induced TNF-α and increased IL-10, indicating a direct, cell-intrinsic effect on macrophage polarization rather than a consequence of reduced recruitment alone. These observations are consistent with prior work establishing LPA as a chemotactic and pro-inflammatory mediator for myeloid cells (49), including our own prior findings that autotaxin inhibition reduces myocardial inflammation after ischemic injury (15–17). These findings are relevant to HFpEF as previous evidence reinforces that monocyte-derived and resident cardiac macrophages actively shape the diastolic phenotype based on the facts that CCR2+ monocyte-derived macrophages drive early hypertrophic remodeling (50), resident macrophages restrain fibrosis and support angiogenesis (51), and macrophage-mediated inflammation links cardiomyocyte oxidative stress to diastolic dysfunction (52). Altered immune cell signatures, both resident and circulating, are now recognized as a conserved feature of HFpEF across species (53, 54), and by lowering the ATX/LPA activity that sustains this deranged myeloid response, autotaxin inhibition addresses a proximal driver rather than a downstream consequence. Upon further exploration of the potential mechanism underlying the anti-inflammatory role of PF8380, it was evident that PF8380 modulates the anti-inflammatory role by preventing cytoplasmic-to-nuclear translocation of NF-κB in mouse bone marrow-derived cells in an in vitro model, thereby reducing TNF-α expression and concomitantly inducing the anti-inflammatory IL-10 and IL-6. Although IL-6 exhibits both pro-inflammatory and anti-inflammatory roles in different pathological conditions, and is usually elevated in the HFpEF condition (55), it is worth noting that IL-6 deficiency leads to adult-onset obesity in mouse (56), and hepatic IL-6 signaling suppresses hepatic inflammation and improves systemic insulin action (57), and drives macrophage polarization from pro-inflammatory to anti-inflammatory phenotype (58, 59).

Beyond inflammation, PF8380 reduced two structural determinants of diastolic dysfunction, interstitial fibrosis and cardiomyocyte hypertrophy, and improved echocardiographic indices of relaxation (E/A ratio and isovolumic relaxation time). A pro-fibrotic role for the autotaxin-LPA axis is established across organs, including the lung (60), kidney (28), and heart (13), and LPA receptor 1 antagonism has advanced to clinical testing in idiopathic pulmonary fibrosis and reverses steatohepatitis in preclinical models (61). Our data extend this anti-fibrotic principle to HFpEF and link it to functional improvement. Whether the diastolic benefit reflects reduced fibrotic stiffness, improved cardiac energetics, or both cannot be resolved from the present data, but the convergence with mitochondrial and metabolic mechanisms recently implicated in cardiometabolic HFpEF (62, 63) suggests these pathways are not mutually exclusive.

These findings have therapeutic implications. The pharmacologic therapies with demonstrated benefit in HFpEF, sodium-glucose cotransporter 2 inhibitors and incretin-based agents, act principally on the metabolic substrate, and sodium-glucose cotransporter 2 inhibition improves diastolic function in part through mitochondrial mechanisms (63). Autotaxin inhibition is mechanistically distinct, targeting the inflammatory and fibrotic sequelae of the metabolic milieu rather than the metabolic derangement itself. In our model, PF8380 improved diastolic function and exercise capacity without a large change in body composition, indicating that its benefit is not simply a consequence of weight loss. This raises the possibility of a complementary strategy in which autotaxin inhibition is combined with metabolic agents to address the inflammation and remodeling that persist despite adequate metabolic control.

Several limitations warrant consideration. First, PF8380 is a pharmacologic inhibitor, and off-target contributions cannot be fully excluded. Second, the elevation of circulating autotaxin in patients is cross-sectional and does not establish a temporal or causal relationship to disease progression; LPA signaling is transduced through at least six receptors with context-dependent and occasionally opposing actions, so circulating enzyme or ligand levels may not track linearly with tissue-level signaling. Third, although BayesPrism deconvolution mitigates the cell-type averaging inherent to bulk RNA sequencing, cell-type-specific and spatial confirmation, ideally by single-nucleus sequencing, is needed to validate the immune and fibroblast programs we infer. Finally, the two-hit murine model, although it reproduces the metabolic, hypertensive, and diastolic features of HFpEF, does not capture the full heterogeneity of the human syndrome, and translation will require testing in additional models and, ultimately, in clinical studies.

In conclusion, we demonstrate that the autotaxin-LPA axis is activated in HFpEF and that pharmacologic inhibition of this axis reduces systemic inflammation, cardiac myeloid cell accumulation, fibrosis, and hypertrophy, while improving diastolic function and exercise capacity. By coupling human lipidomic and inflammatory profiling with mechanistic interrogation of a murine model across the transcriptomic, cellular, and functional levels, these findings identify autotaxin as a therapeutic target that addresses the inflammatory and fibrotic core of HFpEF. Whether autotaxin inhibition can complement existing metabolic therapies to alter the natural history of this common and lethal syndrome is a question that now merits clinical investigation.

## Acknowledgments.

Dr. Abdel-Latif is supported by the VA Merit award (I01CX002684-01), NIH R61/R33 (1R61HL177474-01), and the Mathers Foundation grant.

## Ethics approval

All animal procedures were approved by the University of Michigan Institutional Animal Care and Use Committee and were conducted in accordance with the Guide for the Care and Use of Laboratory Animals, and the study is reported in accordance with the ARRIVE 2.0 guidelines.

---

**Supplementary Figure 1.**
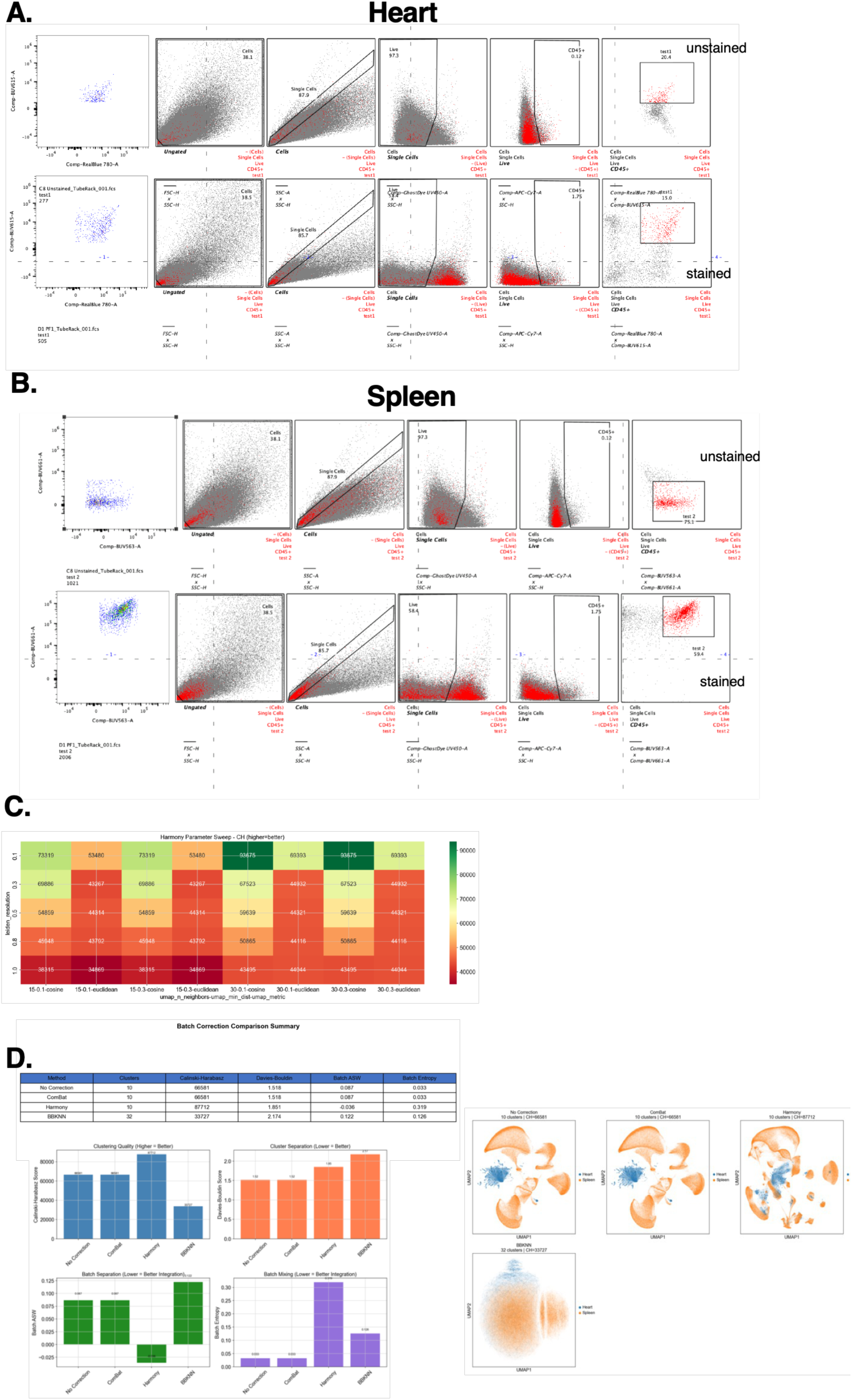
Data preprocessing, parameter optimization, and batch correction for CAFE analysis of spectral flow cytometry data. **(A)** FlowJo pre-gating strategy for cardiac tissue. Unstained control (top row) and representative stained sample (bottom row) are shown side by side. The gating hierarchy proceeds from an initial biaxial plot (Comp-BUV615-A vs Comp-RealBlue 780-A) through cell identification (FSC-H vs SSC-H), singlet discrimination (SSC-A vs SSC-H), viability gating (Ghost Dye UV450 vs SSC-H), CD45+ leukocyte selection (APC-Cy7 vs SSC-H), and a final marker gate (Comp-RealBlue 780-A vs Comp-BUV615-A). Unstained cardiac tissue demonstrated minimal CD45+ events (0.12%), confirming the specificity of the CD45 gate in the presence of cardiomyocyte autofluorescence. Gates were adjusted to minimize inclusion of autofluorescent non-immune cells. **(B)** FlowJo pre-gating strategy for spleenic tissue. Unstained control (top row) and representative stained sample (bottom row) showing the same sequential gating hierarchy as in (A), with initial biaxial plot using Comp-BUV661-A vs Comp-BUV563-A. The higher CD45+ frequency in spleen compared to heart reflects the expected enrichment of leukocytes in lymphoid tissue. **(C)** Harmony parameter optimization. Heatmap of Calinski-Harabasz scores (higher values indicate better clustering quality) across combinations of Leiden clustering resolution (0.1–1.0), UMAP n_neighbors (15, 30), min_dist (0.1, 0.3), and distance metric (cosine, Euclidean). Final parameters selected: resolution = 0.5, n_neighbors = 30, min_dist = 0.1, cosine metric. **(D)** Comparison of batch correction methods for integration of heart and spleen spectral flow cytometry data. Summary table and bar charts comparing four approaches — No Correction, ComBat, Harmony, and BBKNN — across four metrics: Calinski-Harabasz score (clustering quality; higher is better), Davies-Bouldin score (cluster separation; lower is better), batch average silhouette width (Batch ASW; lower indicates better tissue integration), and batch entropy (lower indicates better batch mixing). Corresponding UMAP visualizations are colored by tissue of origin (heart, blue; spleen, orange) for each method. Harmony provided optimal batch integration while preserving biological cluster structure and was selected for all downstream analyses.

**Supplementary Figure 2.**
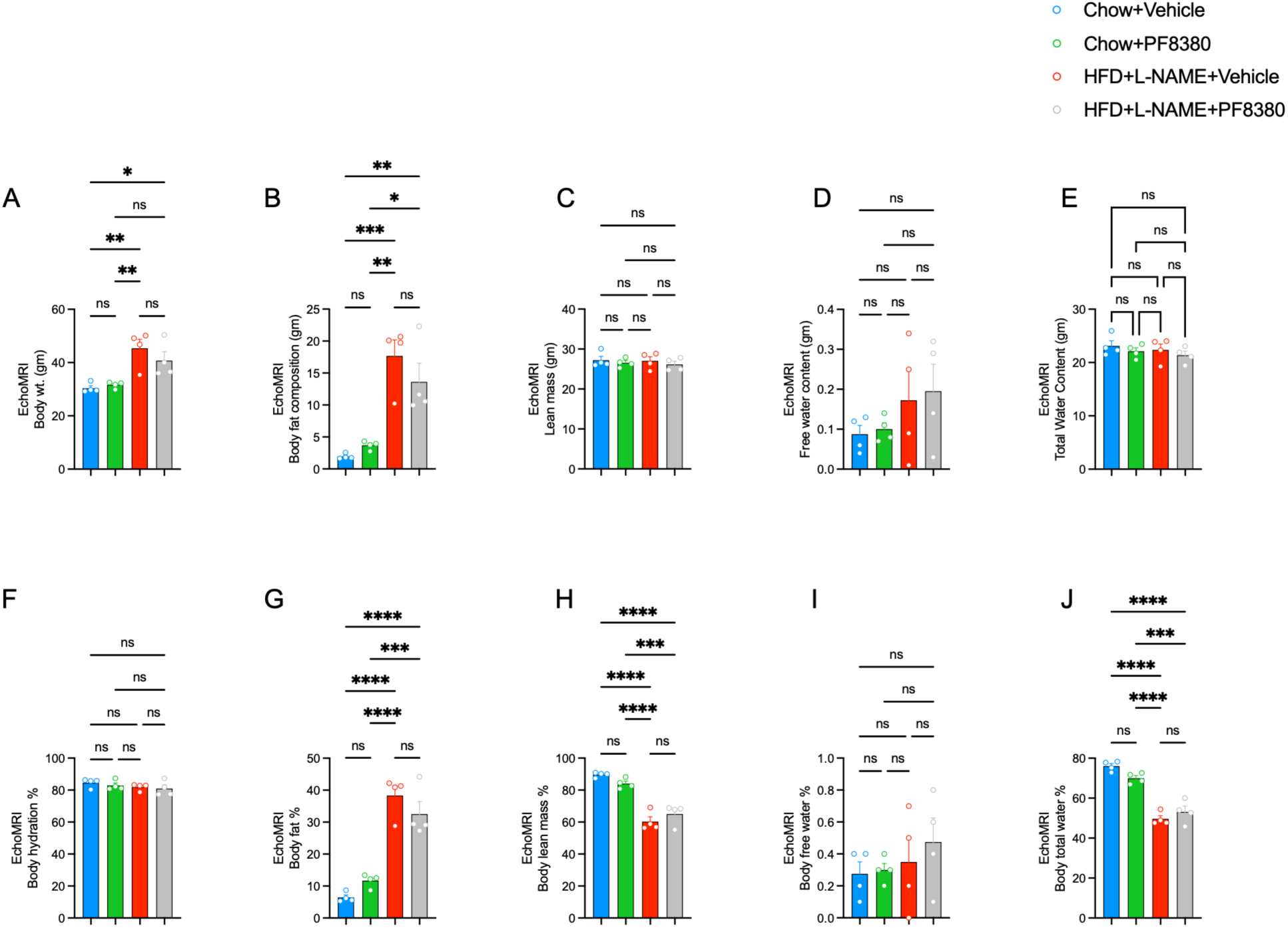
EchoMRI-mediated body composition measurements at week-15. **(A-E)** Body weight, body fat composition, lean mass, free water content, and total water content in grams (n=3-4/group). **(F-J)** Percentage body hydration, body fat, body lean mass, body free water, and body total water (n=4/group). **EchoMRI:** Echo magnetic resonance imaging. **gm:** grams**, HFD:** high-fat diet, **L-NAME:** N-nitro L-arginine methyl ester. **%:** percentage. One way ANOVA with Tukey’s posthoc test. *P<0.05, **P<0.01, ***P<0.001, and ****P<0.0001.

**Supplementary Table 1.**

| **\*Cytek Aurora spectral flow panel - Innate immune cell panel** |  |  |  |  |  |
| --- | --- | --- | --- | --- | --- |
|  | **Antibody** | **Color** | **Cells**<br>**(Spleen)/beads** | **Amount of ab** | **Company/cat#** |
| 1 | Unstained |  | Cells/beads | n/a | n/a |
| 2 | CD115<br>(CSF-1R) | BV421 | Beads | 2.5uL (1uL control) | Biolegend #135513 |
| 3 | CD11b | BUV615 | Beads | 1.25uL (1uL control) | BD #751140 |
| 4 | CD11c | BV750 | Beads | 1 uL | Biolegend #117357 |
| 5 | mCCR2 | BUV661 | Cells | 3uL | BD 750042 |
| 6 | CD45 | APC-Cy7 | Cells/beads | 1.25uL (0.2uL control) | Biolegend #103116 |
| 7 | CD68/SR-D1 | BV711 | Beads | 1.25uL (1uL control) | Biolegend #137029 |
| 8 | CD86 | BV480 | Beads | 1.25uL (1uL control) | BD 746370 |
| 9 | F4/80 | BUV563 | Cells/Beads | 1.25uL (1uL control) | BD 749284 |
| 10 | Ly6G | RB780 | Beads | 1.25uL (0.5uL control) | BD 569152 |
| 11 | Ly-6G/Ly-6C (Gr-1) | SB780 | Beads | 1.25uL (0.5uL control) | eBioscience #78-5931-82 |
| 12 | LYVE1 | AF488 | Beads | 1.25uL (0.2uL control) | eBioscience #53-0443-82 |
| 13 | Timd4 | PE | Beads | 1.25uL (0.2uL control) | eBioscience #12-5866-82 |
| 14 | CD64 | PE-Cy5 | Beads | 1uL (0.2uL control) | Biolegend #139332 |
| 15 | MHC-II | eF506 | Cells/beads | 1uL (0.2uL control) | eBioscience #69-5321-82 |
| 16 | CD161 (NK1.1) | BV605 | Beads | 5uL (1uL control) | Biolegend #108740 |
| 17 | Live Dead | Ghost Dye UV450 | Cells/ViaComp Beads | 1uL (0.1uL control beads) | Cytek Biosciences #13-0868 |

